# Efficacy of glucocorticoid modulator PT150 as a weight loss strategy

**DOI:** 10.64898/2026.04.06.712688

**Authors:** Victoria Glass, Molly McDougle, Whitnei Smith, Parker Dhillon, Lanah Ha, Jose H. Ledo, Christopher D. Verrico, Estefania P. Azevedo

## Abstract

Obesity affects millions of people worldwide and has serious complications such as cardiovascular disease and diabetes. Current treatments for obesity target proteins such as the receptors for glucagon-like peptide-1 (GLP-1), gastric inhibitory polypeptide (GIP) and/or glucagon (GCG). These interventions have revolutionized the treatment of obesity and represent first-line pharmacotherapeutic strategies. One major weakness to these strategies is that once drug treatment stops, most patients are unable to maintain the new body weight setpoint, often gaining weight back rapidly. Thus, the identification of new therapies that focus on the ability to maintain homeostatic setpoint are necessary. The glucocorticoid receptor (GR) has been implicated in several pathways including reward-seeking, inflammation, stress and energy balance. Here, we investigated the effects of 30 days treatment with PT150 (40 mg/kg), a novel GR antagonist, alone and in combination with semaglutide (30 nmol/kg) on food intake, glucose homeostasis, body weight and setpoint maintenance using a C57Bl/6 diet-induced obesity (DIO) mouse model. We monitored food intake and body weight throughout treatment and after drug washout for 20 days to evaluate defended body weight maintenance (body weight setpoint). Our results indicate that treatment with PT150 alone does not significantly alter body weight but in combination with semaglutide it shows the most promising effects in body weight reduction and homeostatic setpoint maintenance. Together, these data suggest that PT150, a GR modulator, may be effective as a homeostatic setpoint modulator when combined with semaglutide.

## Introduction

Obesity is a chronic disease that affects hundreds of millions of individuals worldwide and significantly increases the risk of cardiovascular disorders, type 2 diabetes, and even certain types of cancers (1). In recent years, pharmacological therapies targeting incretin signaling such as glucagon-like peptide-1 receptor (GLP1R) agonists and dual or triple GLP-1/GIP/GCG receptor agonists, have revolutionized the treatment of obesity, increasing weight loss in both preclinical models and clinical populations (2,3). Despite their efficacy, a limitation of these therapies is their inability to maintain a reduced body weight (a new, lower homeostatic setpoint) once treatment is discontinued (4).

In addition to challenges in weight maintenance, current anti-obesity drugs are associated with side effects that limit tolerability and long-term adherence (5,6). Many patients require chronic treatment to sustain weight loss, further increasing the burden of adverse effects (5,6). Together, these limitations highlight a critical gap in the field: the need for therapeutic strategies that not only induce weight loss but also stabilize the post-weight-loss homeostatic setpoint, thereby reducing reliance on continuous drug exposure.

The regulation of body weight is strongly influenced by stress-responsive neuroendocrine systems, among which glucocorticoid signaling plays a central role (7). Glucocorticoids act through the glucocorticoid receptor (GR) to regulate energy balance, feeding behavior, adiposity, and metabolic adaptation during both physiological and pathological states (7). Dysregulated glucocorticoid signaling has been linked to obesity, visceral fat accumulation, insulin resistance, and altered stress-related feeding (7). Conversely, modulation of GR signaling has been shown to influence food intake, reward-related behaviors, and body weight regulation in both preclinical and clinical contexts (8-10). A good example is mifepristone, a GR antagonist for which weight loss has been reported in certain studies (11). However, mifepristone also acts on other endocrine pathways, including progesterone receptor signaling (8), limiting its suitability as a safe and selective therapeutic strategy for obesity. Importantly, GR signaling operates at the intersection of central stress circuits and peripheral metabolic tissues, positioning it as a compelling target for interventions aimed at long-term energy balance rather than acute appetite suppression.

PT150 is a novel glucocorticoid receptor antagonist, with reduced affinity for progesterone receptors relative to mifepristone, that has been shown to attenuate glucocorticoid-dependent behavioral and physiological outcomes in preclinical models (12-14). Prior work has demonstrated that PT150 reduces stress-associated behaviors and alters glucocorticoid-driven phenotypes without the broad endocrine disruption observed with classical GR antagonists (12-14). These properties raise the possibility that PT150 could modify stress-dependent drivers of weight regain and homeostatic setpoint defense, particularly when used in combination with agents that induce robust initial weight loss, such as GLP1R agonists.

In the present study, we tested the hypothesis that targeting GR signaling with PT150 could enhance the durability of weight loss induced by semaglutide, a clinically relevant GLP1R agonist (15). Using a diet-induced obesity (DIO) mouse model, we examined the effects of PT150 alone and in combination with semaglutide on body weight, food intake, glucose tolerance, and pancreatic immune cell infiltration during active treatment and following drug washout. By explicitly incorporating a washout period, our study was designed to assess not only weight loss efficacy but also the capacity of these interventions to stabilize defended body weight setpoint after treatment cessation. Our findings demonstrate that while semaglutide drives robust weight loss during active treatment, only the combination of PT150 and semaglutide confers protection against post-treatment weight regain. Importantly, this effect occurs without persistent changes in food intake, suggesting that GR modulation may engage mechanisms distinct from classical appetite suppression. Together, this work provides a conceptual and experimental framework for developing adjunct therapies aimed at maintaining weight loss by targeting glucocorticoid signaling pathways, addressing a major unmet need in obesity treatment and offering new avenues for improving the long-term efficacy of current pharmacotherapies.

## Materials and Methods

### Animals

C57BL/6J DIO male mice aged between 21-24 weeks were purchased from Jackson Laboratories and used throughout this study. Mice were allowed access to food and water *ad libitum* and were housed on a 12 hr light/dark cycle. All animal experiments were approved by the MUSC IACUC following the National Institutes of Health guidelines for the Care and Use of Laboratory Animals. Mice were put on 60% high fat diet (HFD) (Research Diets Cat#D12492) at 6 weeks of age for 15 weeks (DIO mice were purchased from Jackson Laboratories at a final age of 21-24 weeks). At 15 weeks of HFD exposure, animals were given drug treatments accordingly and remained on HFD until the end of the study (end of drug washout period at week 22).

### Drug Treatment

At week 15 (21-24 weeks of age), animals received PT150 (40 mg/kg/daily) administered orally in 1g of Nutella and semaglutide (30 nmol/kg/daily) administered i.p. in sterile saline following previously standardized protocols (15,16). Vehicle controls received both saline injections and 1g of Nutella daily for 30 days or 4 weeks (week 15-19). Drugs and vehicle controls were prepared fresh daily. After drug treatment and during drug washout, no vehicle or drugs were administered for an additional 20 days (week 19-22). PT150 was a gift from Palisades Therapeutics and Dr. Mark Prendergast at University of Kentucky. Semaglutide was obtained from Sigma/AA Blocks.

### Glucose Tolerance Test

At the end of the drug washout period (week 22), animals were fasted overnight for 14-16hrs and then given an i.p. bolus of 1.5g/kg of glucose (Sigma). Blood glucose was tested using a glucometer (AccuCheck) at 0, 15, 30, 60 and 120 min. After experiments, mice were given food *ad libitum* and returned to their home cage.

### Food Intake and Body Weight Monitoring

To facilitate food measurement, animals were single housed 1 week before (starting at week 14) and during the remainder of the experiment until euthanasia. Before and during the 30 days of drug administration and washout, food intake and body weight were measured every 3-4 days manually by a blind experimenter using a commercial scale.

### Immunohistochemistry

At the end of experiments (week 22), mice were perfused with 1X PBS and 10% normal buffered formalin (NBF), and pancreas were postfixed for 24 hr in 10% NBF. Pancreas were cryoprotected in 30% sucrose for 24hrs each or until organs sunk. Tissue was then embedded in OCT and stored at -80C until slicing. Tissue slices were taken using a cryostat (Leica), blocked for 1 hr with 0.3% Triton X-100, 3% bovine serum albumin (BSA), and 2% normal goat serum (NGS) and incubated in primary antibodies for 24 hr at 4°C on a shaker. Then, slices were washed three times for 10 min in 0.1% Triton X-100 in PBS (PBS-T), incubated for 1 hr at room temperature with secondary antibodies, washed in PBS-T and mounted in Vectamount with DAPI (Southern Biotech). Antibodies used here were anti-CD68 (1:500; Thermo Scientific) and goat-anti-rat (Alexa 594, 1:1000). Images were taken using AxioImager.M2 microscope (Zeiss) and images were processed and cells counted using ImageJ software (NIH).

### Analyses

All results are presented as mean ± SEM and were analyzed with Graphpad Prism software 9.0 (GraphPad). Unless otherwise indicated, we used a one-way ANOVA with Bonferroni correction to analyze statistical differences. p<0.05 was considered significant (*p<0.05, **p<0.01, ***p<0.001, ****p<0.0001. Different treatment groups were blinded to investigators in the various experiments.

## Results

### PT150 in combination with semaglutide slows body weight regain after drug treatment cessation

To assess the effects of PT150, semaglutide, and their combination on body weight regulation in diet-induced obese (DIO) mice given a 60% HFD, animals were treated for 4 weeks (30 days) followed by a 20-day drug washout period (Fig. 1A). Body weight was monitored throughout the experiment (Fig.1B and C), and body weight changes (Δ in g; Fig.1C) were calculated relative to baseline (day 0). During the treatment phase, vehicle-treated mice exhibited a progressive increase in body weight over time as expected (black symbols and lines; 17). In contrast, semaglutide treatment (blue symbols and lines) induced a rapid and significant reduction in body weight, reaching an average of 10g of body weight loss at the end of the 4-week treatment period (***p<0.001, vehicle vs semaglutide groups; Fig.1B-D). PT150 alone (red symbols and lines) significantly prevented weight gain when compared to vehicle (**p<0.01, Fig.1B-D). Notably, combined PT150 and semaglutide treatment (purple symbols and lines) resulted in a significant weight loss effect, compared to vehicle controls and similar to semaglutide alone during the treatment phase, with a robust reduction in body weight relative to vehicle (with an average of 10g of weight loss; Fig.1B-D). Analysis of body weight change at the end of drug treatment revealed a significant body weight loss in the PT150+semaglutide group compared with vehicle (***p <0.001; Fig.1D).

**Figure 1.**
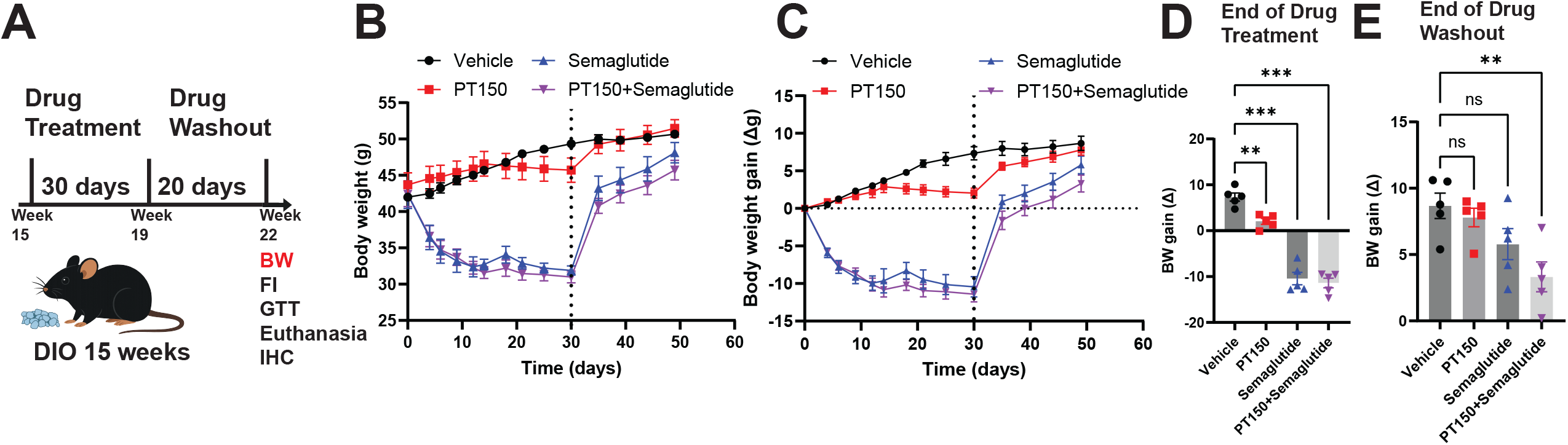
PT150 combined with semaglutide slows body weight regain after treatment cessation in DIO mice. DIO mice (15 weeks on diet-induced obesity regimen) received 30 days or 4 weeks of treatment with vehicle (black symbols), PT150 (red symbols), semaglutide (blue symbols), or PT150+semaglutide (purple symbols), followed by 20 days of drug washout. **(A)** Experimental timeline illustrating treatment and washout phases and associated outcome measures (BW, FI, GTT, Euthanasia and tissue collection for IHC). **(B)** Longitudinal body weight (g) over time. The dotted vertical line marks the transition from drug treatment to washout. **(C)** Longitudinal body-weight change (Δ in g) over time. **(D)** Quantification of body weight change at the end of drug treatment and **(E)** end of washout. Each symbol represents one mouse; bars represent mean ± SEM. N=5 mice males. One-way ANOVA with Bonferroni post hoc test was performed as shown: **p < 0.01, ***p < 0.001; ns, not significant.

Following drug washout, all groups exhibited partial rebound in body weight, seen as early as day 35 (5-days after drug washout). However, differences between treatment groups persisted. At the end of the washout period, PT150+ semaglutide-treated mice were the only group that displayed significantly lower body weight gain compared with vehicle (**p<0.01; Fig.1E). In contrast, PT150 alone and semaglutide alone were not significantly different from vehicle after washout. This shows that although semaglutide had significant effects in weight loss, these effects were not sustained. Together, these data indicate that semaglutide drives the dominant weight-reducing effect during active treatment, while PT150 alone has limited efficacy on body weight in DIO mice. However, only combination treatment confers significant protection against post-treatment weight regain, maintaining the defended body weight setpoint in mice.

To determine whether differences in body weight trajectories were driven by sustained changes in food intake, cumulative food intake (FI) was measured every 3-4 days throughout the treatment and washout phases in diet-induced obese (DIO) mice (Fig. 2A). Although semaglutide-treated and PT150+semaglutide-treated mice showed a significant reduction in cumulative FI relative to vehicle during treatment (Fig.2B and C, ***p<0.001), these differences were transient and did not reach statistical significance at the end of our experiment (Fig.2B and D). PT150 alone showed no significant effects in the first days of treatment (Fig.2B and C). At the end of drug treatment (day 30), PT150 alone also did not significantly alter cumulative FI when compared with vehicle at any time point (Fig.2B and D). During the 20-day washout period, cumulative FI increased similarly across all groups, with evidence of modest, but non-significant, compensatory hyperphagia in the semaglutide treated groups of mice (Fig.2E). Together, these data indicate that in our experiments neither PT150, semaglutide, nor their combination produces sustained alterations in cumulative food intake, suggesting that the body weight effects observed with PT150+semaglutide are not due to a suppression in food intake.

**Figure 2.**
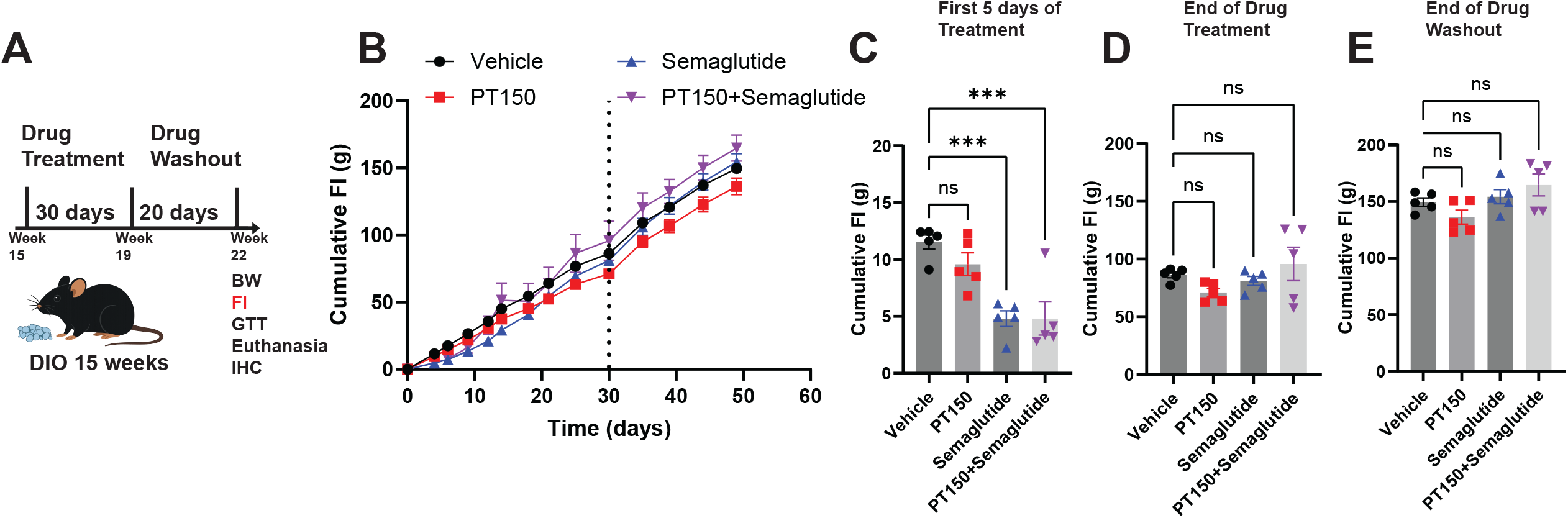
PT150 does not produce sustained changes in cumulative food intake in DIO mice. DIO mice (15 weeks on diet-induced obesity regimen) received 4 weeks of treatment with vehicle (black symbols), PT150 (red symbols), semaglutide (blue symbols), or PT150+semaglutide (purple symbols), followed by 20 days of drug washout. **(A)** Experimental timeline illustrating treatment and washout phases and associated outcome measures (BW, FI, GTT, Euthanasia and tissue collection for IHC). **(B)** Cumulative food intake (g) measured longitudinally across the study. The dotted vertical line denotes the transition from drug treatment to washout. **(C)** Cumulative food intake at the first 5 days of drug treatment. **(D)** Cumulative food intake at the end of drug treatment. **(E)** Cumulative food intake at the end of drug washout. Each symbol represents an individual mouse; bars indicate mean ± SEM. N=5 male mice. One-way ANOVA with Bonferroni post hoc test was performed as shown: ns, not significant.

### PT150 Improves Glucose Tolerance Alone and in Combination with Semaglutide

To determine whether chronic PT150 treatment alters glucose handling in diet-induced obese (DIO) mice fed a 60% HFD, animals were treated for four weeks with vehicle, PT150, semaglutide, or PT150 combined with semaglutide, followed by a 20-day drug washout prior to glucose tolerance test (GTT; Fig. 3A). All treatments showed a significant reduction in the area under the curve suggesting appropriate disposal of glucose, and thus a recovery from glucose resistance (*p<0.05, Fig.3B and C). Peak glucose occurred between 15-30 min post-injection across groups, with similar glucose trajectories during the clearance phase across treated groups (Fig.3B). These data indicate that all treatments produce a persistent alteration in glucose tolerance after cessation of treatment under these experimental conditions.

**Figure 3.**
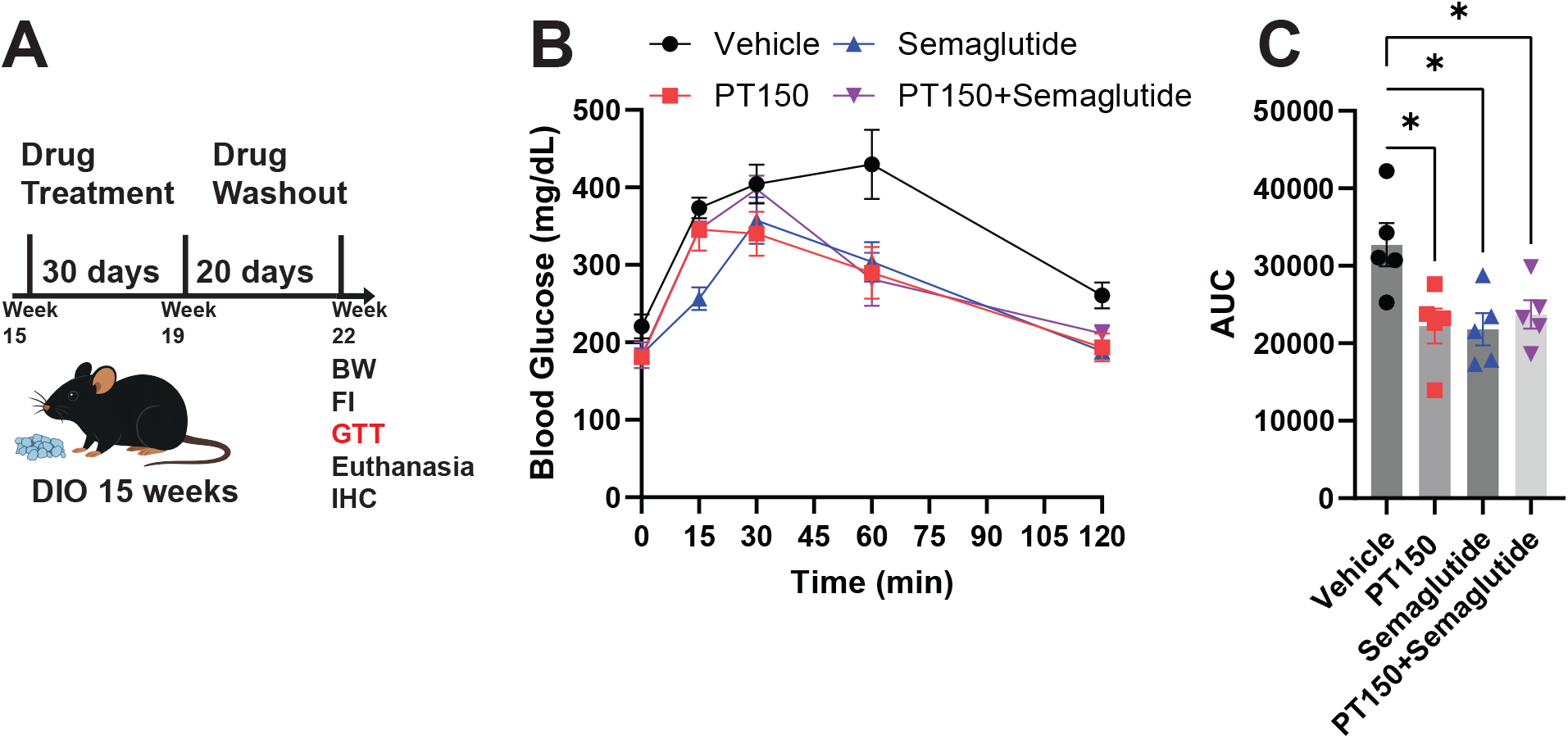
Effects of PT150 on glucose tolerance following drug washout in DIO mice. DIO mice (15 weeks on diet-induced obesity regimen) received 4 weeks of treatment with vehicle (black symbols), PT150 (red symbols), semaglutide (blue symbols), or PT150+semaglutide (purple symbols), followed by 20 days of drug washout. **(A)** Experimental timeline illustrating treatment and washout phases and associated outcome measures (BW, FI, GTT, Euthanasia and tissue collection for IHC). **(B)** Blood glucose levels (mg/dl) measured during an intraperitoneal glucose tolerance test (1.5 g/kg i.p. bolus) performed at the end of the washout period. **(C)** Quantification of glucose area under the curve (AUC). Each symbol represents an individual mouse; bars indicate mean ± SEM. N = 5 male mice per group. Statistical comparisons were performed using one-way ANOVA with Bonferroni post hoc tests as indicated; *p < 0.05.

### PT150 Does Not Produce Persistent Changes in Pancreatic Immune Cell Density Following Drug Washout

To evaluate whether chronic PT150 treatment produces sustained changes in pancreatic immune cell composition in diet-induced obese (DIO) mice fed a 60% HFD, pancreatic tissue was collected following a 20-day drug washout period after chronic treatment with vehicle, PT150, semaglutide, or PT150 combined with semaglutide (Fig. 4A). Pancreatic sections were analyzed by immunohistochemistry for CD68 (red, Fig.4B), a marker of macrophages/monocytes, to assess treatment-related alterations in myeloid cell abundance. Quantification of CD68-positive (CD68+) cell density did not differ between vehicle-treated mice and mice previously treated with PT150, semaglutide, or the combination therapy (Fig. 4B and C). These findings indicate that neither PT150 alone nor in combination with semaglutide induces persistent changes in pancreatic macrophage cell number following treatment cessation under the conditions tested.

**Figure 4.**
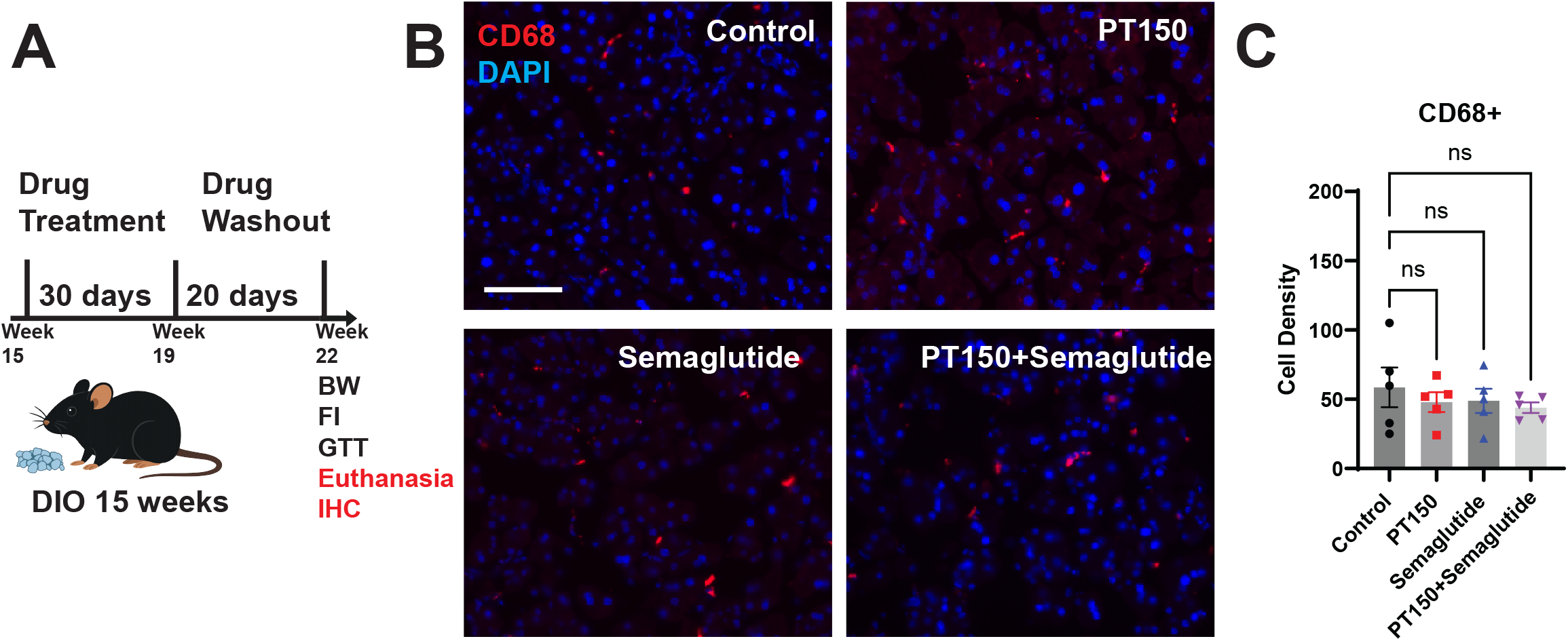
Analysis of the effects of PT150 on pancreatic immune cell number following drug washout in DIO mice. DIO mice (15 weeks on diet-induced obesity regimen) received 4 weeks of treatment with vehicle (black symbols), PT150 (red symbols), semaglutide (blue symbols), or PT150+semaglutide (purple symbols), followed by 20 days of drug washout. **(A)** Experimental timeline illustrating treatment and washout phases and associated outcome measures (BW, FI, GTT, Euthanasia and tissue collection for IHC). **(B)** Representative immunofluorescence images of pancreatic sections from each treatment group stained with DAPI (blue) and CD68 (red). **(C)** Quantification of CD68-positive (CD68^+^) cell density in pancreatic tissue at the end of the washout period. Immunopositive cell density expressed as cells per image. Each symbol represents an individual mouse; bars indicate mean ± SEM. Scale bars, 100 μm. N = 5 male mice per group. Statistical comparisons were performed using one-way ANOVA with Bonferroni post hoc tests as indicated; ns, not significant.

## Discussion

In this study, we examined the effects of the glucocorticoid receptor antagonist PT150, alone and in combination with the GLP1R agonist semaglutide, on body weight regulation, food intake, glucose homeostasis, and pancreatic immune cell composition in a diet-induced obesity (DIO) mouse model. Our findings demonstrate that while semaglutide drives the dominant body weight loss effects, as expected during active treatment, the combination of PT150 and semaglutide uniquely confers durable protection against post-treatment weight regain following drug washout (Fig.1). Importantly, this sustained effect occurs in the absence of persistent changes in cumulative food intake (Fig.2) or pancreatic immune cell infiltration (Fig.4). Additionally, PT150 treatment alone or in combination with semaglutide improved glucose tolerance (Fig.3), suggesting that PT150 may modulate body weight setpoint through mechanisms distinct from acute appetite suppression.

In this study, although we observed a significant decrease in food intake in the first 5 days of treatment with semaglutide or PT150+semaglutide groups, no robust differences in cumulative food intake across treatment conditions were observed at the end of drug treatment (day 30). These findings are consistent with prior reports (15) which demonstrated only transient, early reductions in food intake at a 30 nmol/kg (∼2g/day reduction in intake at day 5 after treatment started) dose following semaglutide treatment for 21 days (15).

Our study also analyzed the ability of PT150 alone or in combination with semaglutide to improve glucose tolerance in DIO mice, which at >15 weeks showed glucose intolerance (Fig. 3). Following a 20-day drug washout, glucose tolerance was improved, as expected, in the semaglutide groups but also in the PT150 alone or the PT150+semaglutide combinatorial group. This was not unexpected as PT150 functions as a GR antagonist, which has been associated with improvements in insulin sensitivity and glucose tolerance (11)

A concern for an increased risk of pancreatitis has been raised for many weight loss drugs, including semaglutide (5). Based on that reasoning, our study conducted pancreatic histological analyses after 20 days of drug washout to assess whether immune cell density to the pancreas occurred. Our study showed that the sustained weight maintenance observed with PT150 + semaglutide was not associated with persistent alterations in pancreatic immune cell infiltration (CD68, a macrophage marker). Another important clinical concern for weight loss treatments such as semaglutide is increased gastric emptying and gastroparesis (5,18). The current study did not perform studies to assess gut motility or gastric emptying in treated mice, but we believe that future studies aiming to better define safety in the combination of PT150+ semaglutide treatment will be needed, including the assessment of changes in gut function.

Finally, our study suggests that a combinatorial treatment of PT150+semaglutide decrease rapid body weight regain after chronic semaglutide treatment. Whether these effects can be expanded to other weight loss drugs remain to be studied. Additionally, the mechanisms by which PT150 sustain weight loss in combination with semaglutide remain to be defined. One intriguing possibility is that PT150 may alter gastrointestinal physiology or nutrient handling, thus changing the physiological satiety signals. Another possibility is that PT150 may act centrally to modulate stress-responsive or homeostatic circuits that govern defended body weight, potentially stabilizing the post-weight loss setpoint established by semaglutide treatment.

### Study Limitations

In our study, we acknowledge the existence of limitations that should be explored in future work. Here we tested a single dose of semaglutide and PT150 and therefore did not evaluate dose-response relationships or alternative dosing strategies that may influence weight loss magnitude or durability. Our experiments focused exclusively on semaglutide, and it remains unknown whether similar protective effects on weight regain would be observed when PT150 is combined with dual or triple incretin receptor agonists such as tirzepatide or GLP-1/GIP/glucagon tri-agonists. Glucocorticoid signaling, GLP1R responsiveness, and energy balance regulation are all known to exhibit sex differences (18-20). Only male mice were included in this study, thus, future studies with larger cohorts of both sexes designed to explicitly examine sex as a biological variable will be critical to determine whether PT150 exerts sexually dimorphic effects on energy balance or body weight setpoint regulation. In addition, the post-treatment follow-up period was limited to 20 days, preventing conclusions about longer-term weight regain dynamics and sustained setpoint stabilization. Finally, mechanistic studies were beyond the scope of the present work, and the central and peripheral pathways through which PT150 enhances durability of weight loss remain to be defined.

In summary, our data identify PT150 as a promising adjunct to GLP1R agonist therapy, capable of enhancing the durability of weight loss after treatment cessation without exacerbating glucose intolerance or inducing pancreatic inflammation. Although the precise mechanisms remain unresolved, these findings serve as framework and support the concept that targeting GR signaling may complement existing anti-obesity therapies by stabilizing body weight setpoint rather than solely suppressing food intake.

## Acknowledgements

We thank the veterinarian staff at DLAR at MUSC for technical assistance. We thank Palisades Therapeutics and Dr. Prendergast at University of Kentucky for providing PT150. We thank Dr. Mehboob Hussain at UCI Health for important manuscript suggestions.

## Author Contributions

E.P.A. designed the study. E.P.A., M.M., W.S., V.G., P.D. performed and analyzed experiments. L.H. provided technical assistance with experiments. E.P.A, J.L, C.D.V. and V.G. wrote the manuscript with input from all authors.

## Data Availability

Further information and requests for reagents may be directed to, and will be fulfilled by the corresponding author Estefania Azevedo.

